# Novel Vβ specific germline contacts shape an elite controller T cell response

**DOI:** 10.1101/2020.05.31.102566

**Authors:** Yang Wang, Alexandra Tsitsiklis, Wei Gao, H. Hamlet Chu, Yan Zhang, Wei Li, Wing Ki Wong, Charlotte M. Deane, David Neau, Jill E. Slansky, Paul G. Thomas, Ellen A. Robey, Shaodong Dai

## Abstract

Certain CD8 T cell responses are particularly effective at controlling infection, as exemplified by elite control of HIV in individuals harboring HLA-B57. To understand the structural features that contribute to CD8 T cell elite control, we focused on a strongly protective CD8 T cell response directed against a parasite-derived peptide (HF10) presented by an atypical MHC-I molecule, H-2L^d^. This response exhibits a focused TCR repertoire dominated by Vβ2, and a representative TCR (TG6) in complex with L^d^-HF10 reveals an unusual structure in which both MHC and TCR contribute extensively to peptide specificity, along with a parallel footprint of TCR on its pMHC ligand. The parallel footprint is a common feature of Vβ2-containing TCRs and correlates with an unusual Vα-Vβ interface, CDR loop conformations, and Vβ2-specific germline contacts with peptide. Vβ2 and L^d^ may represent “specialist” components for antigen recognition that allow for particularly strong and focused T cell responses.

## Introduction

A key strategy for adaptive immune recognition in mammals is to generate enormous diversity by somatic rearrangements of antigen receptor genes during T and B cell development. While B cells can realize the full potential of this diversity, allowing them to recognize virtually any molecular structure, T cell recognition is highly constrained by the requirement for presentation of antigenic peptides by Major Histocompatibility Complex (MHC) proteins, and the need for the T cell antigen receptor (TCR) to recognize both the antigenic peptide and polymorphic self-MHC proteins. How T cells achieve both broad coverage and high specificity, given the constraints imposed by MHC restriction, is a central question in T cell biology.

Part of the answer to this question may come from the binding orientation of the TCR on its peptide MHC (pMHC) ligand^1^. In the vast majority of known TCR-pMHC structures, the TCR docks in a diagonal orientation, such that the highly variable complementarity determining (CDR) 3 loops, which are encoded by somatic rearrangement joints, are positioned directly over the peptide, the most variable part of the pMHC ligand. In contrast, CDRs 1 and 2, which are germline-encoded within individual variable (V) gene segments, contact the MHC α-helices that make up the sides of the peptide binding groove. There is evidence that tyrosine residues within the TCR CDR1 and 2 loops help to impose this characteristic docking angle, although this remains controversial, and random selection models have also been proposed^2, 3, 4^. Due to the large number of allelic forms of MHC, the impact of CDR3, and the flexible geometry of TCR-pMHC interactions, identifying conserved germline contacts requires comparing multiple TCR-pMHC structures. Indeed, most of our current understanding of conserved germline contacts comes from mouse Vβ8 containing TCRs, and related Vβs in human, which contain tyrosine residues within their CDR1 and 2s, and are the most represented TCR-pMHC structures in the Protein Data Bank^2, 5, 6^. It remains unclear whether TCRs that use divergent Vβs segments have a similar docking orientation and germline contacts with pMHC.

In addition to the TCR docking orientation, the peptide binding characteristics of MHC also contribute to the broadness and specificity of antigen recognition. Peptides generally bind in an extended conformation within a deep groove of MHC, and the ability to bind to distinct allelic forms of MHC is largely determined by 2 or 3 peptide “anchor” residues. This arrangement ensures that peptide binding is sufficiently broad that a handful of MHC molecules in an individual can present peptides from virtually any pathogen, with the fine specificity for peptide determined largely by the TCR. On the other hand, there are indications that atypical MHC-I molecules, with relatively restricted peptide binding, can nevertheless generate strong CD8 T cell responses. For example, the ability of certain individuals to control HIV infection without anti-retroviral therapy, termed “elite control”, is associated with HLA-B alleles (e.g. B27 and B57) with limited ability to bind peptides^7, 8^. Restricted peptide binding by mouse H2-L^d^ is correlated with resistance to CD8 T cell exhaustion during chronic infection^9^, and is controlled by amino acid polymorphisms that also correlate strongly with HIV control^7, 10^. The paradoxical association between restricted peptide binding and elite control may be due to the combination of weak binding to self-peptides coupled with strong binding to particular antigenic peptides^8, 9^. However, precisely why certain MHC-I molecules favor the development of particularly potent CD8 T cell responses remain a mystery.

The potent T cell response to the intracellular protozoan parasite, *Toxoplasma gondii* in resistant (H2^d^) mouse strains is dominated by CD8 T cells specific for a single peptide, HF10, (derived from the parasite protein GRA6) presented by L^d11^. The L^d^-HF10 specific T cell response exhibits a number of similarities to CD8 T cell responses in HIV elite controller patients, including the lack of T cell exhaustion in the face of persistent infection and continuous production of armed effector T cells via a proliferative intermediate T cell population^12, 13, 14^. This unusually potent T response may serve as a model for understanding CD8 T cell responses that underlie strong resistance to viral infection in certain individuals.

Here we show that the L^d^-HF10-specific T cell response displays a focused TCR repertoire dominated by Vβ2. Crystallographic studies of a representative TCR (TG6) in complex with L^d^-HF10 reveal an unusual parallel footprint on the pMHC complex, a feature that is also observed in other Vβ2 containing TCRs, and which promotes germline-encoded TCR contacts with bound peptide. In addition, the HF10 peptide binds tightly and with high complementary to L^d^, in a conformation that optimizes peptide side chain interactions with both the MHC and TCR. Thus this example of T cell elite control uses a strategy in which both TCR and MHC contribute substantially to peptide specificity, and runs counter to the prevailing view of T cell recognition. We discuss these results in terms of a model in which both Vβ2 and MHC-I L^d^ represent “specialist” recognition components that sacrifice broad coverage in order to provide unusually strong and focused responses to particular pathogens^15^.

## Results

### The L^d^-HF10 specific T cell response is characterized by early activation, strong expansion, and a focused TCR repertoire

*T. gondii* infection of genetically resistant (H-2^d^) mice elicits a potent CD8 T cell response directed against a single parasite-derived peptide, associated with the continuous production of armed effector T cells during chronic infection^11, 12^. To examine the priming of this T cell response in more detail, we performed pMHC tetramer staining of splenic T cells during the 2 weeks following i.p. infection of F1(B6xB6.C) mice. Consistent with previous results, T cells specific for the GRA6 derived 10mer peptide HF10 presented by MHC-I L^d^ expanded >10^4^ fold, compared to a <100x expansion by the subdominant responses (Fig. 1a). The strong expansion of L^d^-HF10 specific T cells corresponds to a greater up-regulation of activation and effector markers at day 5 post infection compared to subdominant T cell responses (Fig. 1b). A similar expansion and immunodominance hierarchy of *T. gondii* epitopes was observed following oral infection (Supplementary Fig. 1).

**Fig. 1.**
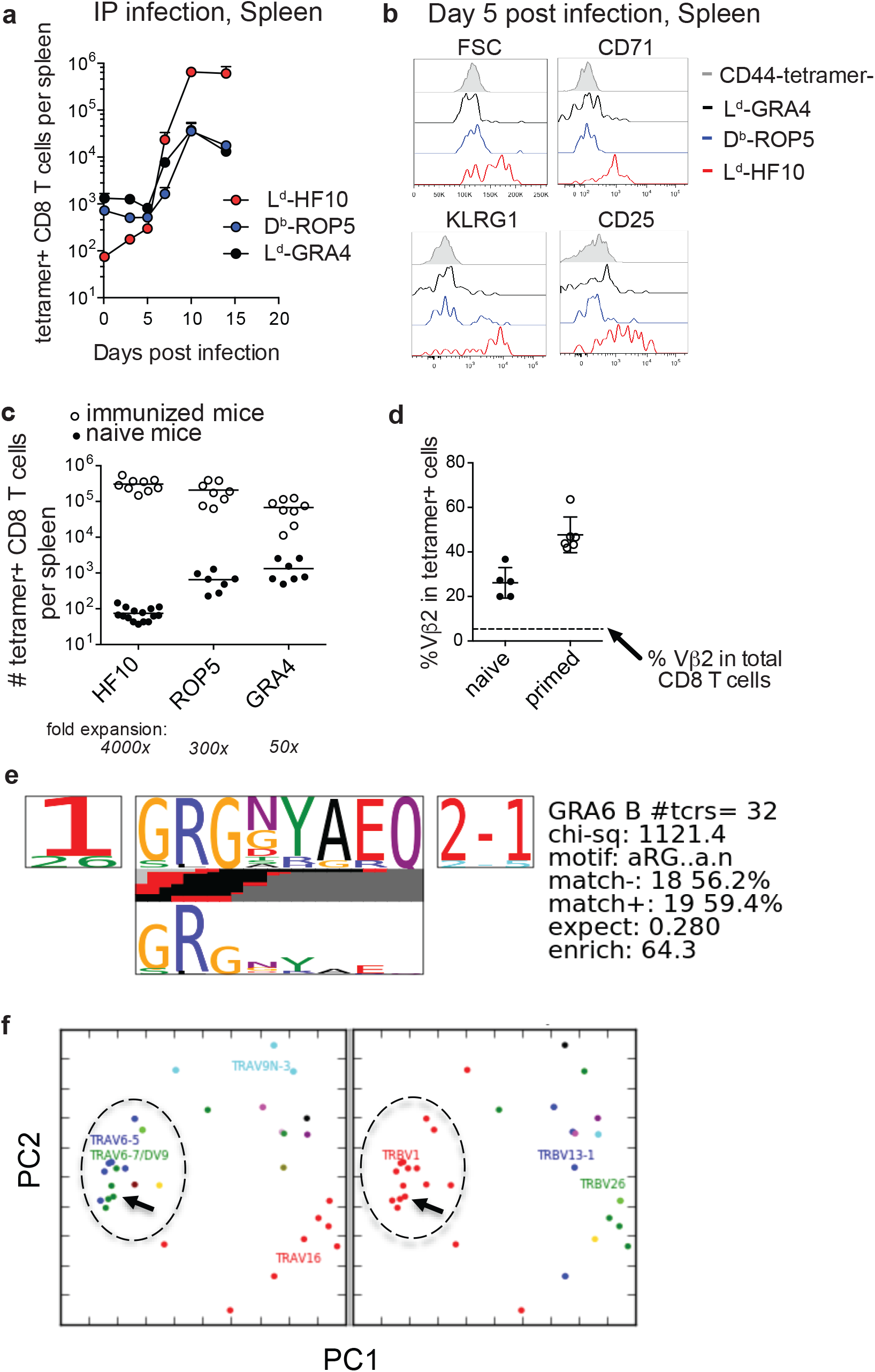
Characteristics of the L ^d^-HF10 specific T cell response. **a**, *T. gondii*-specific CD8 T cells were quantified by pMHC tetramer staining and flow cytometry of splenocytes at different time points after intraperitoneal infection of F1 (B6xB6.C) mice. **b**, Flow cytometric analysis of size (FSC or forward scatter) or expression of activation and effector markers (CD71, KLRG1, and CD25) on gated tetramer^+^ splenic CD8 T cells at day 5 post infection. **c**, Mice were immunized with bone marrow-derived dendritic cells loaded with the indicated peptides. Expanded tetramer^+^ CD8 T cells were quantified by tetramer staining of splenocytes 7 days post immunization (open circles). Numbers of tetramer^+^ CD8 T cells in the spleen were quantified by tetramer enrichment of naïve mice and were used to calculate the fold expansion of each antigen-specific T cell population (closed circles). **d**, The frequency of Vβ2 usage amongst L^d^-HF10 specific splenic CD8 T cells tetramer enriched from naïve mice or found in *T. gondii*-infected mice was determined by flow cytometry. Each dot represents an individual mouse and the dashed line indicates the frequency of Vβ2 amongst total splenic CD8 T cells (5.40%). **e**-**f**, L^d^-HF10 tetramer^+^ CD8 T cells were sorted from mice 3 weeks post infection and TCRα and TCRβ genes from individual T cells were sequenced as described^48^. Clonal diversity in L^d^-HF10 specific CD8 T cells was analyzed using the TCRdist algorithm^18^. **e**, Top-scoring CDR3β motif. Results of a CDR3 motif discovery algorithm are shown using a TCR logo that summarizes V and J usage, CDR3 amino acid enrichment, and inferred rearrangement structures. The bottom panel shows the motif enriched by calculating against a background dataset of non-epitope specific TCR sequences. **f**, Principal components analysis (PCA) projection of the TCRdist landscape colored by Vα (left panel) and Vβ (right panel) gene usage. The groups of TCRs that correspond to the top scoring CDR3β motif are indicated with a dashed circle, and the TG6 TCR is indicated with an arrow.

Previous studies have shown that the secretion pattern of the antigenic precursor protein and the C-terminal location of the HF10 epitope contribute to, but do not fully account for, the strong L^d^-HF10 response^16, 17^. To determine the impact of TCR-pMHC interactions, we examined T cell responses following immunization of naïve mice with peptide-loaded dendritic cells (DC) (Fig. 1c). T cells specific for L^d^-HF10 expanded 4000x upon peptide-DC immunization, whereas T cells specific for the other epitopes showed substantially lower expansion (Fig. 1c). These data indicate that the interactions between TCR, peptide, and MHC, as well as parasite biology and antigen presentation, contribute to the potency of the L^d^-HF10 T cell response.

We previously demonstrated that L^d^-HF10 specific T cells from infected mice show preferential usage of Vβ2^11^. This preference for Vβ2 is also observed upon immunization of mice with DC loaded with the HF10 peptide, whereas T cell responses to other peptides displayed a Vβ profile that closely matched that of bulk CD8 T cells (data not shown, and Supplementary Fig. 1). L^d^-HF10 specific T cells from naïve mice also showed a Vβ2 frequency significantly above that of bulk CD8 T cells (Fig. 1d) suggesting that a preference for Vβ2 is already evident after thymic selection, and increases after T cell priming with the HF10 antigenic peptide.

To further investigate the TCR repertoire of the L^d^-HF10 response, we sequenced paired TCR α and β genes from 80 individual T cells from 2 different chronically infected mice, yielding 32 unique paired α/β TCR sequences. We analyzed the unique sequences using the TCRdist algorithm^18^. The majority of T cells used the TRBV1 segment (which encodes Vβ2) together with TRBJ2-1 and displayed a strong selection for a GRG motif in the TCRβ CDR3 (Fig. 1e). We also noted a preference for TCRβ CDR3 length of 13 amino acids, a trend that was particularly prominent for Vβ2 containing T cells (Supplementary Fig 1). Finally, principal component analysis based on TCR distances showed a predominant cluster of similar TCRs which included the previously identified TG6 TCR (Fig. 1f)^12^. Indeed, the TG6 TCRβ coding sequence from the original L^d^-HF10 specific T cell hybridoma was found independently in the 2 additional mice examined, each time paired with a closely related TCRα. Likewise, the TCRα gene of the TG6 TCR was found independently in one additional mouse, paired with a closely related TCRβ gene (Supplementary Table 1). These data indicate that the response to L^d^-HF10 displays a highly focused TCR repertoire dominated by Vβ2. Moreover, TG6 is a Vβ2-containing TCR that provides a good representative of this response.

### Unusual peptide conformation and non-anchor contacts characterize HF10 binding to L^d^

Previous studies revealed unusual features of L^d^, including a constrained peptide binding site and the requirement for non-anchor residues for optimal peptide binding^10, 19, 20^. To characterize the binding of the HF10 peptide to L^d^, we expressed recombinant soluble L^d^ molecules containing a covalently linked HF10 peptide and solved the crystal structure at 1.8 Å resolution. As described for other L^d^-bound peptides, HF10 does not lie flat, but instead bends within the peptide-binding groove to accommodate an obstruction formed by aromatic stacking interactions between elite-control associated residues including 97W in L^d^ (Fig. 2a, Supplemental Video 1)^8, 10^. However, while previously described L^d^-bound peptides (all 9mers) have a bend at either p5 or p6 of the peptide^10, 20, 21, 22^, HF10 displays pronounced bends at both p5 and p7 (Fig. 2b and Supplementary Fig. 2). Thus, the extra length of the HF10 peptide is accommodated by small bends in the peptide and a close fit with MHC, without the pronounced bulge outside of the MHC that is often observed with longer than optimal peptides.

**Fig. 2.**
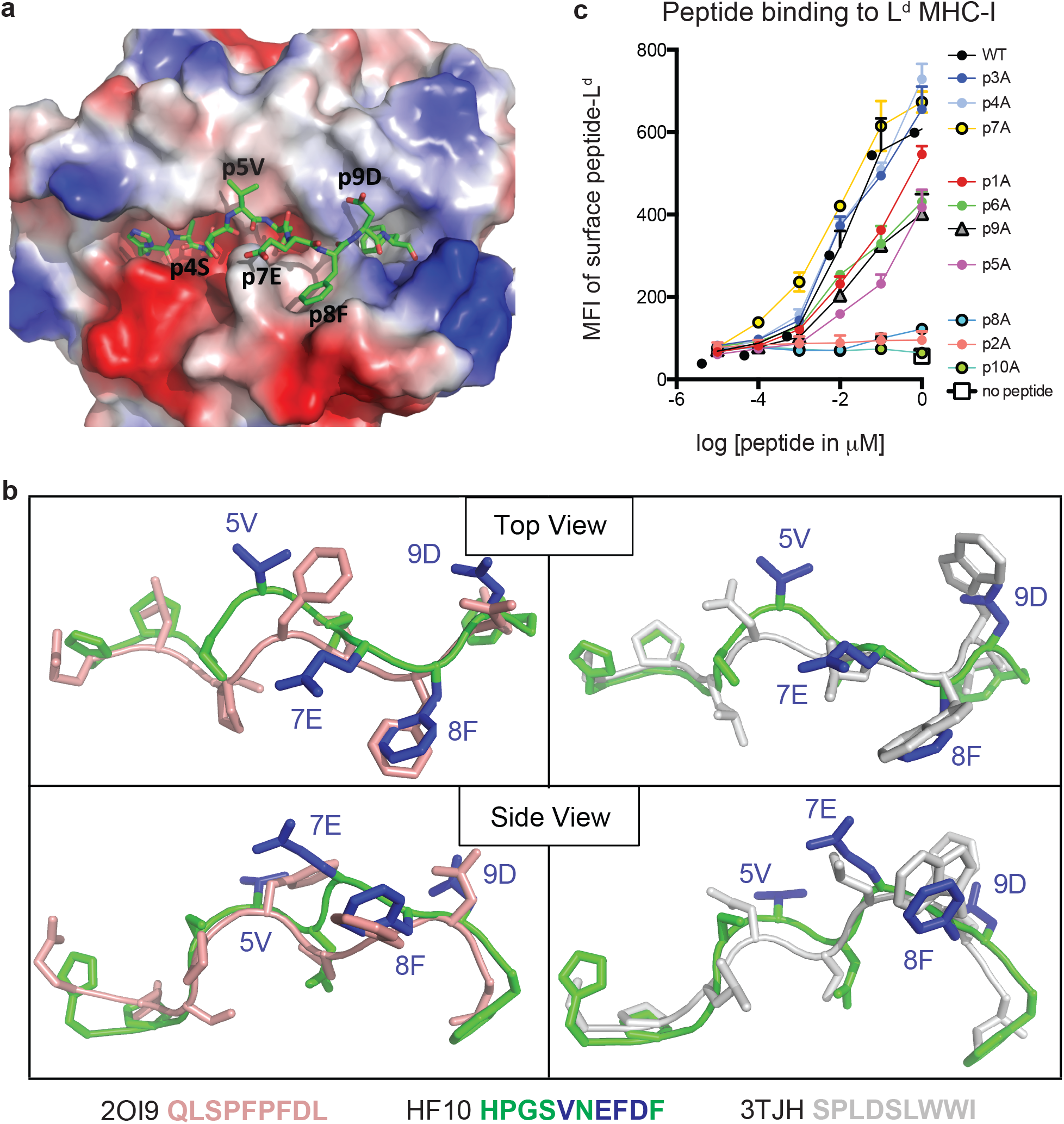
Features of the antigenic HF10 peptide bound to H2-L^d^. Soluble H2-L^d^ containing a covalently linked HF10 antigen peptide was crystallized and the structure was solved at 1.8 Å (Methods and Supplementary Table 2). **a**, L^d^ is shown as an electrostatic potential molecular surface with the HF10 peptide shown in green. Solvent exposed peptide residues are labeled. **b**, Conformation of the bound HF10 peptide (green with blue side chains) in comparison to other L^d^-bound peptides. Peptides with a bend at p5 (HF10 and 2OI9 in pink) are on the left, and peptides with a bend at p6 (p7 for HF10) (HF10 and 3TJH in white) are on the right. See Supplemental Video 1. Additional L^d^-bound peptides are shown in Supplementary Fig. 2. **c**, H2-L^d^ binding to HF10 peptide alanine substitution variants. Flow cytometry surface expression (MFI) of L^d^ on TAP-deficient RMA-S.L^d^ cells incubated with increasing concentrations of the indicated HF10 or peptide variants. Data are representative of 3 independent assays.

In line with the published structures of L^d^, p2P and p10F of HF10 are buried in the B and F pockets respectively and serve as anchor residues. The side chain of p6D interacts with the base of the groove, occupying the C pocket. In addition to these buried contacts, residues 4, 5, 7, 8, and 9 project to the sides of the groove (Fig. 2b), with p5V and p9D making extensive contacts with the L^d^ α1 helix, and p4S and p8F contacting the L^d^ α2 helix. Because these peptide side chains project to the sides of the groove, portions of the residues are exposed to solvent, providing potential TCR contacts. Only a single amino acid side chain, that of p7E, is facing away from H2-L^d^, and fully available to engage a TCR (Supplemental Video 1).

To confirm the interactions between HF10 and L^d^, we performed alanine scan mutagenesis of the peptide (Fig. 2c). We measured binding based on the ability of peptides to stabilize surface L^d^ expression on the TAP-deficient RMA-S cell line^17^. As expected, alanine substitution of the two anchor positions, p2P or p10F, abolished peptide binding, as did substitution at position 8. In addition, substitution at positions 1, 5, 6, 9 substantially reduced binding. These data are consistent with the 3D crystal structure and confirm the close complementarity between peptide and MHC molecules, including non-anchor residue contacts.

### An unusual TCR footprint and a dominant peptide contact characterize TG6 TCR interaction with the L^d^-HF10 complex

To investigate the structural features underlying the potent T cell response to Ld-HF10, we solved the crystal structure of the TG6 TCR bound to the L^d^-HF10 complex. We prepared soluble TG6 TCR by expressing and refolding the extracellular domains as described^6^. We used a refolded version of L^d^-HF10 consisting of the α1 and α2 domains with 5 mutations to improve its stability, and with the original tryptophan at position 97 to preserve HF10 peptide binding^22^ (and data not shown). We co-crystallized the refolded L^d^-HF10 and TG6 at a 1:1 ratio and solved the complex structure at a resolution of 2.5Å.

In the majority of reported TCR-pMHC structures, the TCR has a characteristic diagonal docking orientation, in which CDR3 of the TCR α and β chains are positioned over the peptide, CDR1 and 2 of TCRβ are positioned over the MHC α1 helix, and CDR1 and 2 of TCRα are positioned over the α2 helix of MHC-I (or the β1 helix of MHC-II)^1, 2, 23^. In contrast to this consensus, TG6 displays an unusual footprint, in which all the TCRβ CDR loops shift toward the L^d^ α2 helix. As a result, there are 121 TCR contacts to L^d^ α1, but only 23 contacts to L^d^ α2 (Table 1 and Fig. 3a). The buried surface between TG6 and L^d^-HF10 is 1131.8 Å^2^, relatively small compared to the known TCR-pMHC-I complexes^1^.

**Table 1.**
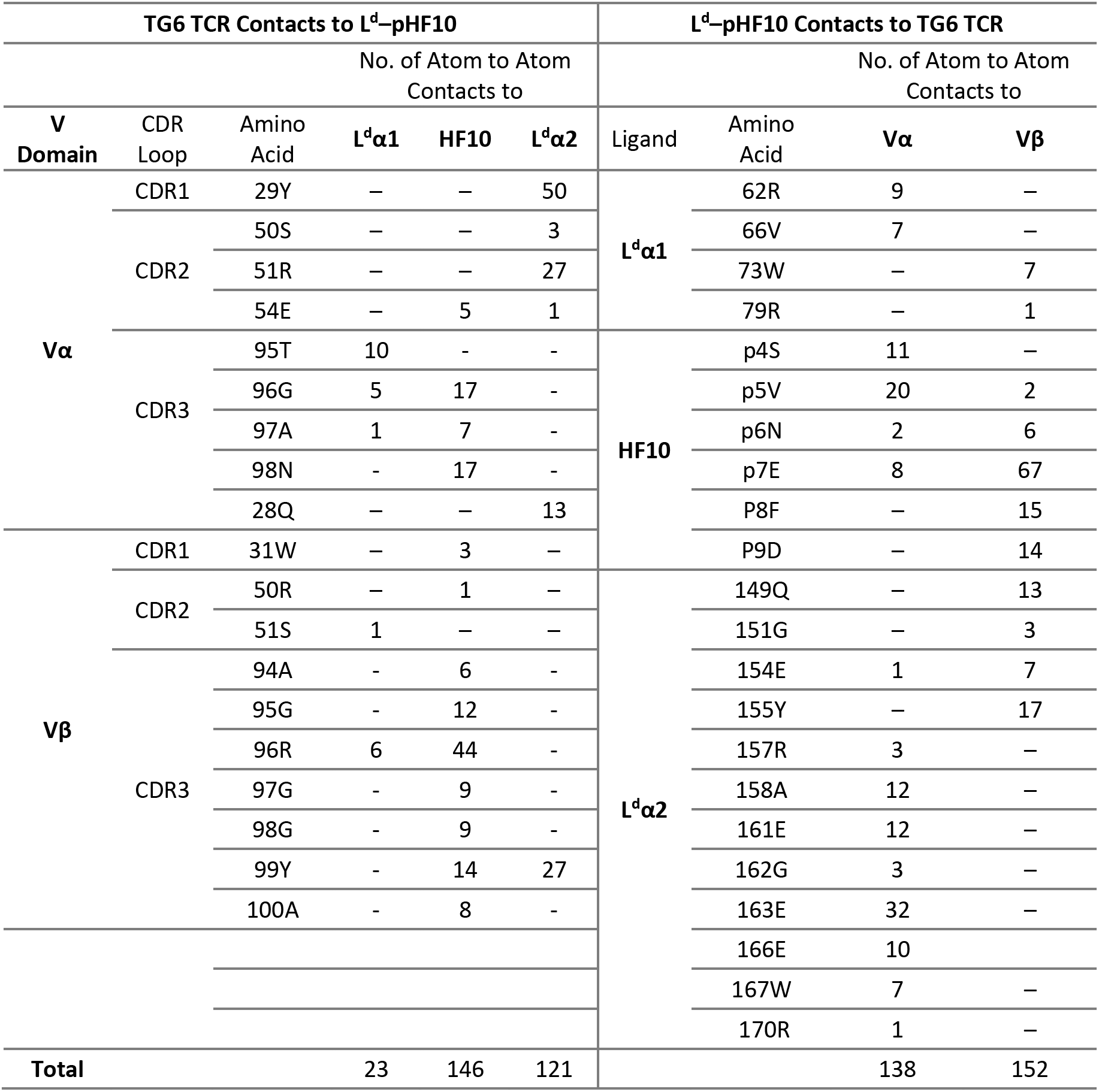
Contacts between TG6 TCR and L^d^–pHF10.

**Fig. 3.**
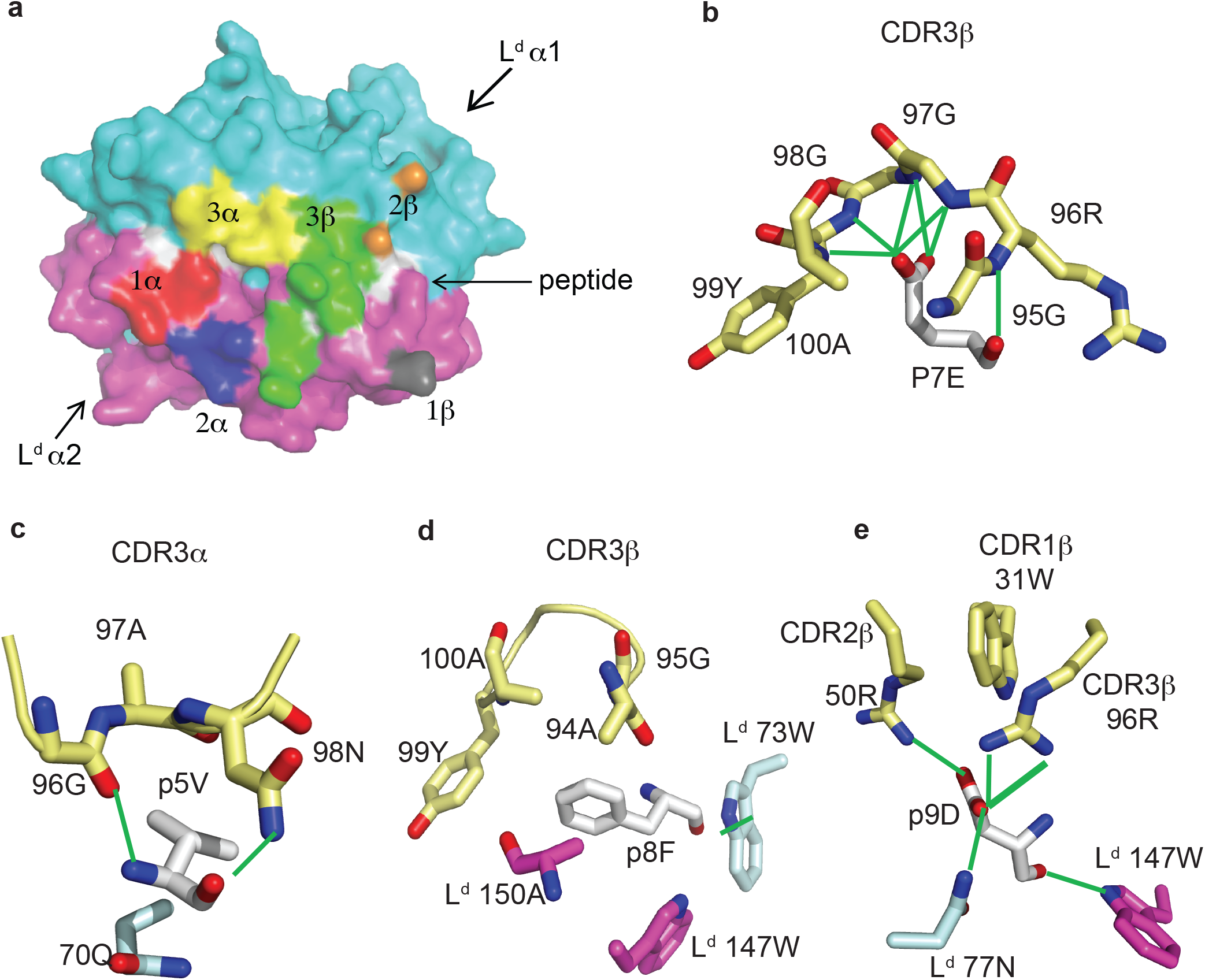
Features of TG6 TCR bound to L^d^-HF10 complex. **a**, TCR footprint on the solvent accessible surfaces of the L^d^-HF10 complexes (L^d^ α1, cyan; L^d^ α2, magenta; peptide, white). Areas of TCR contact with pMHC (≤ 4.5Å) are colored as: CDR1α, red; CDR2α, blue; CDR3α, yellow; CDR1β, gray; CDR2β, orange; CDR3β green. **b-e**, Interactions between TG6 CDRs and L^d^-HF10. HF10 residues are shown in white carbon stick; residues on TG6 are shown in pale yellow carbon stick; residues of L^d^ are shown in magenta (α1) and cyan (α2) carbon stick; H-bonds and salt bridges are indicated by green lines. **b**, Extensive contacts between TCR CDR3β and p7E of the HF10 peptide. **c**, Residues of both TCR CDR3α and L^d^ contact HF10 p5V. **d**, Hydrophobic stacking of TCR CDR3β and L^d^ with HF10 p8F. **e**, Both TG6 CDR2β and CDR3β form a salt bridge to p9D. L^d^ also provides contacts. See Supplemental Video 2.

The TCR interaction with the HF10 peptide is centered around TCRβ chain CDR3 contacts with p7E: the only peptide side chain that points away from the MHC molecule (Fig. 2). This interaction involves a complex hydrogen bond network with the CDR3 backbone (Fig. 3b). The close wrapping of CDR3 loop around p7E is consistent with the conserved length of 13 amino acids, and 2 strongly selected glycine residues in the consensus TCRβ CDR3 determined by TCR sequencing (Fig. 1e, Supplementary Fig. 1, Supplementary Video 2). Additional TCR contacts are formed with peptide residues that project toward the sides of the MHC groove and are sandwiched between the L^d^ α1 and α2 helices and the TCR residues (Fig. 3c-e). These include p5V: which forms hydrogen bond interactions with TG6 CDRα3 via its main chain N and O atoms, as well as van der Waals contacts with its aliphatic side chain (Fig. 3c), p8F: which makes van der Waals contacts with CDR3β (Fig. 3d), and p9D: which is contacted by 96R of CDR3β and 50R from CDR2β (Fig. 3e, Supplementary Video 2). Interestingly, 96R is a prominent part of the enriched motif (GRG) in CDR3 of TCRβ (Fig. 1d), implying that L^d^-HF10 specific TCRs are highly selected to preserve this interaction.

Measurements of TCR binding to pMHC by surface plasmon resonance are in good agreement with the ternary complex structure. The affinity for the wild type peptide is at the high end of the range reported for TCRs (0.4 μM) (Supplementary Fig. 3)^24^. In addition, alanine substitution of p6N, p7E, p8D, and p10F all greatly reduce both TCR binding, and T cell activation (Supplementary Fig. 3). Thus, the C-terminal peptide residues (p6-10) are all crucial for TCR recognition, with p7E contributing exclusively to TCR contacts, and the remaining residues affecting both MHC and TCR interactions (Fig. 2c, Supplementary Fig. 3).

The overlay of the L^d^-HF10 structure before and after engagement of the TG6 TCR shows that, while most of the L^d^-HF10 surface changes little upon TCR binding, there is a shift in the side chain rotamers of L^d^ 155Y and 62R. The side chain of 155Y rotates to point toward the HF10 peptide in the binding groove (Supplementary Fig. 3), allowing the 99Y from TG6 Vβ CDR3 to contact the L^d^ α2 domain via hydrogen bond and van der Waals interactions (Table 1). On the other hand, L^d^ 62R rotates toward the TCR Vα CDR1, allowing the 27S from TG6 to form a hydrogen bond with the L^d^ α2 domain. The large rotamer changes from the L^d^ 155Y and 62R upon TCR binding is consistent with the fit-induced binding kinetics observed in SPR of L^d^-HF10 binding to TG6 (Supplementary Fig. 3), indicating an initial low affinity binding step, followed by a conformational change leading to a more stable complex.

### Vβ2-specific germline contacts correlate with a parallel footprint on pMHC

To understand the basis of the unusual Vβ2 footprint of the TG6 TCR on L^d^-HF10, we compared the structure of TG6 to Yae62, a Vβ8-containing TCR that binds to pMHC with a classic docking orientation and footprint^6^. We noted a shift in the positions of the TCRβ CDR loops of TG6 relative to Yae62, leading to re-positioning of the loops away from the MHC α1 helix, and toward the peptide and MHC α2 helix (Fig. 4a). To explore the basis for the shift in CDR loops, we superimposed the TG6 and Yae62 structures (Fig. 4b, Supplementary Video 3). We noted a striking difference in the conformation of the TCRβ CDR1 and 2 loops, with kinks in the Vβ2 CDR1 and 2 loops due to proline residues at positions 30 and 52. These bends contribute to the re-positioning of the loops toward the peptide and MHC α2 helix. Similar CDR1 and 2 conformations are observed in 3 other Vβ2 containing TCR from 6 different structures (Supplementary Fig. 4).

**Fig. 4.**
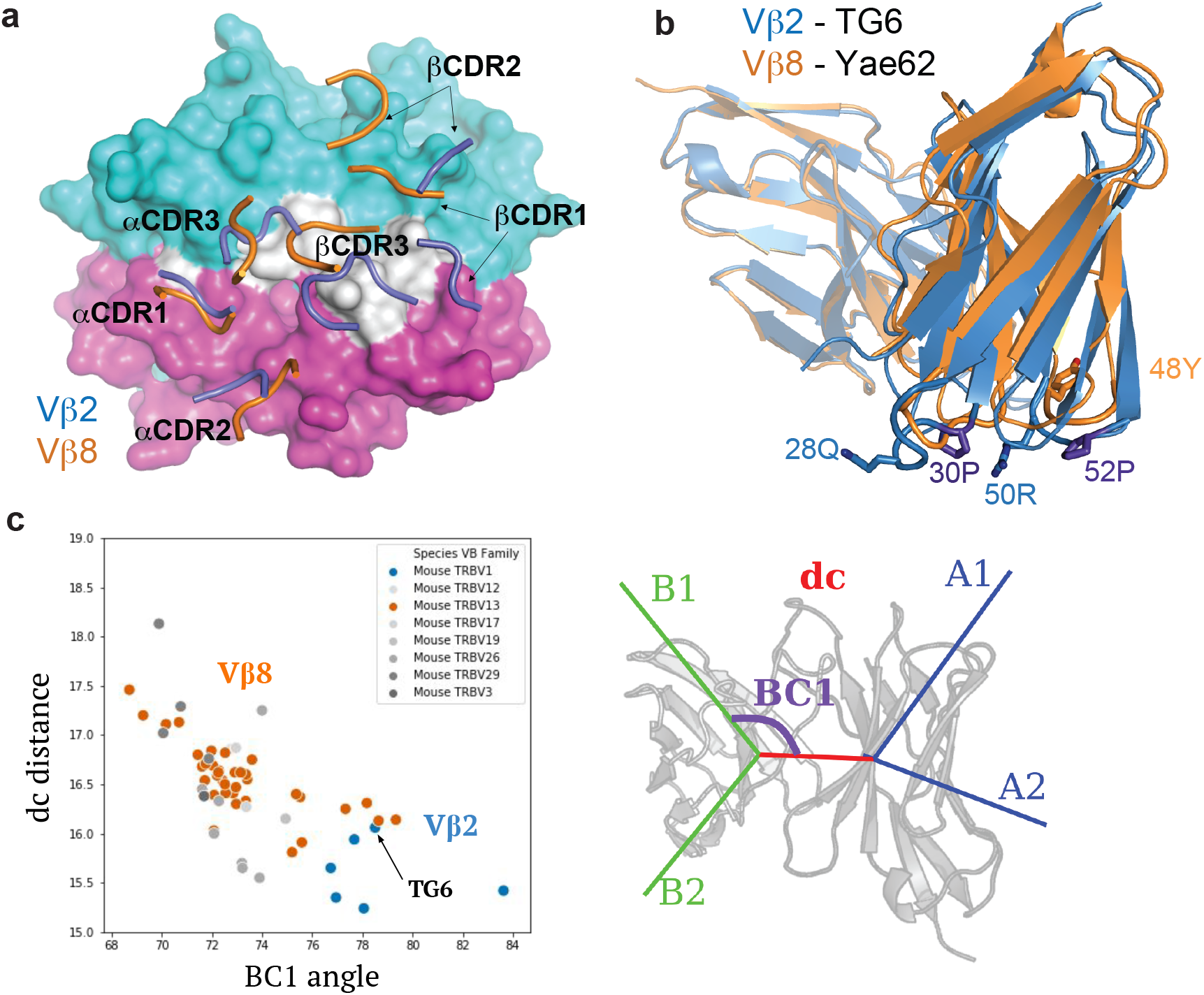
Structural differences between Vβ2 and Vβ8 containing TCRs lead to an altered footprint for pMHC binding. **a**, A comparison of the positions of CDR loops from TG6 and YAe62 TCRs over their pMHC ligands. CDR loops from TG6 are shown in blue and Yae62 are in orange. **b**, Ribbon diagrams of TG6 TCR (Vβ2) and YAe62 TCR (Vβ8) overlaid with their TCRα chains aligned (both Vα4 encoded by TRAV6). Note that there is a shift in the juxtaposition of TCRα and TCRβ domains that contributes to the shift in the position in the CDR loops of TCRα relative to TCRβ. In addition, proline residues in the CDR1 and CDR2 loops of Vβ2 lead to a further shift in the CDR1 and 2 loops away from the α1 helix of MHC and toward the peptide and α 2 helix of MHC-I (or β1 helix of MHC-II) See Supplemental Video 3. .**c**, TRangle parameters dc distance and BC1 angle for the TG6 TCR (indicated by arrow) compared to non-redundant TCR structures in the PDB. Vβ2 TCRs are shown in blue and Vβ8 TCRs are in orange. Right panel shows TRangle parameters used to define the Vα Vβ interface geometry superimposed over a ribbon diagram of TCR.

Another striking difference between TG6 and Yae62 TCRs is the shift in the position of the Vβ domain relative to Vα (Fig. 4b). To quantify the difference in the Vα-Vβ domain interface, we used a method called TRangle, which defines variations in the geometry of Vα-Vβ interface based on 1 distance and 5 angle measurements ^25^. Interestingly, TG6, as well as the other Vβ2 containing TCRs in the Protein Data Bank (https://www.rcsb.org/) have an unusually low DC1 distance, and an unusually high BC1 angle compared to other published mouse TCR structures, most of which use Vβ8 (Fig. 4c). Moreover, these parameters do not show any obvious correlation with the TCRα usage of these same TCRs (Supplemental Fig. 4). Thus both an altered Vα-Vβ interface, and the conformation of the CDR1 and 2 loops contributes to the shift in the TG6 footprint toward the MHC α2 helix.

Given the striking difference between the CDR1/2 loop conformations and Vα-Vβ interface in Vβ2-compared to Vβ8-containing TCRs, we considered that the unusual footprint of TG6 on pMHC might be a common feature with other Vβ2-containing TCRs. Superimposing the TCR footprint of 3 additional Vβ2 TCRs from 5 different structures onto pMHC revealed a similar shift in the TCRβ contacts toward the peptide and MHC α2 helix compared to Vβ8 TCRs (Fig. 5). Typically, TCR docking angles on pMHC are calculated using the positions of conserved V domain cysteines to determine the TCRαβ axis. Since this approach would not capture changes in the Vβ2 footprint due to the unusual CDR2 and 3 conformation, we defined a new parameter that we call the “footprint angle”. First, we selected the CDR residues involved in binding pMHC (using a contact radius of 4.5 angstroms, Table 1). We then defined a TCR vector between the 2 mass centers of the Vα and Vβ CDR contact regions, and calculated the angle between the TCR vector and the peptide vector (defined by the position of the α carbons from residues p1H and p10F). The TCR vectors of all Vβ2 TCRs were relatively parallel to their peptide vectors (footprint angle 14.5 - 27.7 degrees), compared to relatively diagonal TCR vectors for a set of Vβ8 containing TCRs (footprint angle 37 - 53 degrees) (Fig. 5). Thus, the shift in the TCRβ footprint toward a parallel binding orientation on pMHC appears to be a conserved feature of Vβ2 containing TCRs.

**Fig. 5.**
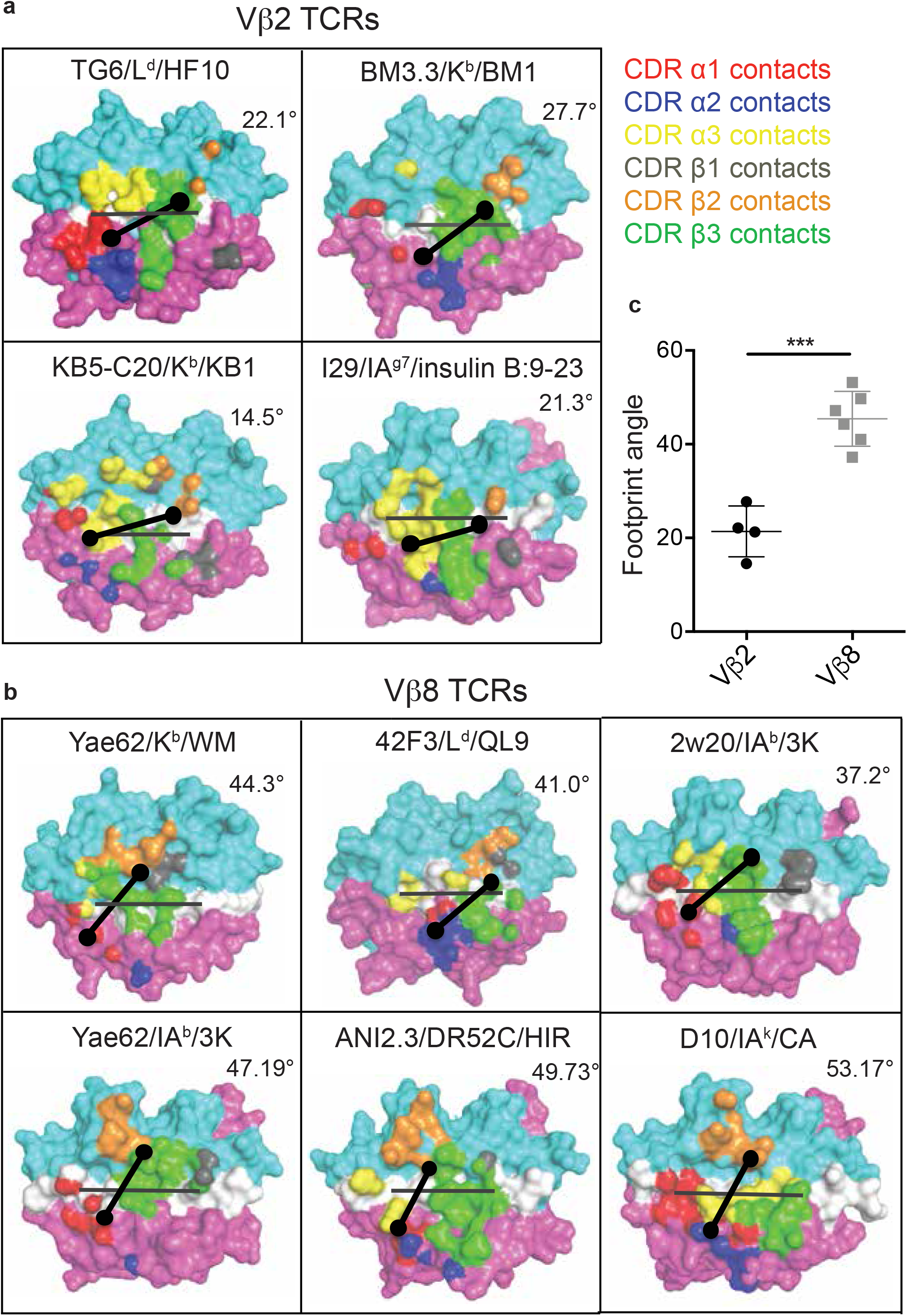
Footprint for binding of Vβ2 and Vβ8 containing TCRs to pMHC. The TCR footprints for 4 different Vβ2 TCR/ pMHC complexes **(a)** and 6 different Vβ8 TCR/pMHC complexes **(b)** are shown, along with footprint angles, calculated based on a vector for the peptide (grey line) and a vector for the TCR-pMHC contact regions (black line) as described in the text. The center of the TCRα and TCRβ footprints are indicated by dots. Structures are: TG6 TCR binding to L^d^-HF10 (PDB: 6X31); BM3.3 TCR binding to K^b^-pBM1 (PDB: 1FO0); KB5-C20 TCR binding to K^b^-pKB1 (PDB: 1JK2); TCR I29 binding to IA^g7^-insulin B:9-23 (PDB: 5JZ4); TCR Yae62 binding to K^b^-pWM (PDB: 3RGV);TCR 42F3 to L^d^-pCPA12 (PDB: 4N5E); TCR 2w20 to IA^b^-3k (PDB: 3C6L); TCR Yae62 to IA^b^-3k (PDB: 3C60); TCR ANI2.3 to DR52c-pHIR (PDB: 4H1L); TCR D10 to IA^k^-pCA (PDB: 1D9K). **c**, Summary of all calculated Vβ2 and Vβ8 TCR footprint angles. Statistical significance was determined by a t-test (***p<0.001).

It has been proposed that the footprint of TCR on pMHC is influenced by germline-encoded contacts, which may differ between particular Vβ segments^2^. To identify conserved Vβ2-specific germline contacts, we compared the interactions of CDR1 and 2 with pMHC in TG6, and 3 other unique Vβ2 containing TCRs (Fig. 6). In all four structures, germline-encoded residue 28Q from the Vβ2 CDR1 contacts the α2 helix of MHC-I or the equivalent β1 helix of MHC-II (Fig. 6a, Table 1). In addition, 50R from the Vβ2 CDR2 contacts both the α1 helix and the peptide in each of the structures. Interestingly, while the aliphatic portion of 50R contacts α76V of MHC-I or α67A of MHC-II, the amino group forms a salt bridge with an acidic residue of the bound peptide in 3 out of 4 of the structures (Fig 6B, Table 1). The previously described Vα germline contact between tyrosine at position 29 with the α2 helix of MHC-I is preserved in TG6 (Supplementary Fig. 5), consistent with the similar TCRα footprint for Vβ2- and Vβ8-containing TCRs on pMHC (Fig. 4a). Thus, Vβ2 specific germline contacts are associated with a shift in the TCRβ footprint that leads to a parallel footprint angle and conserved germline contacts with peptide.

**Fig. 6.**
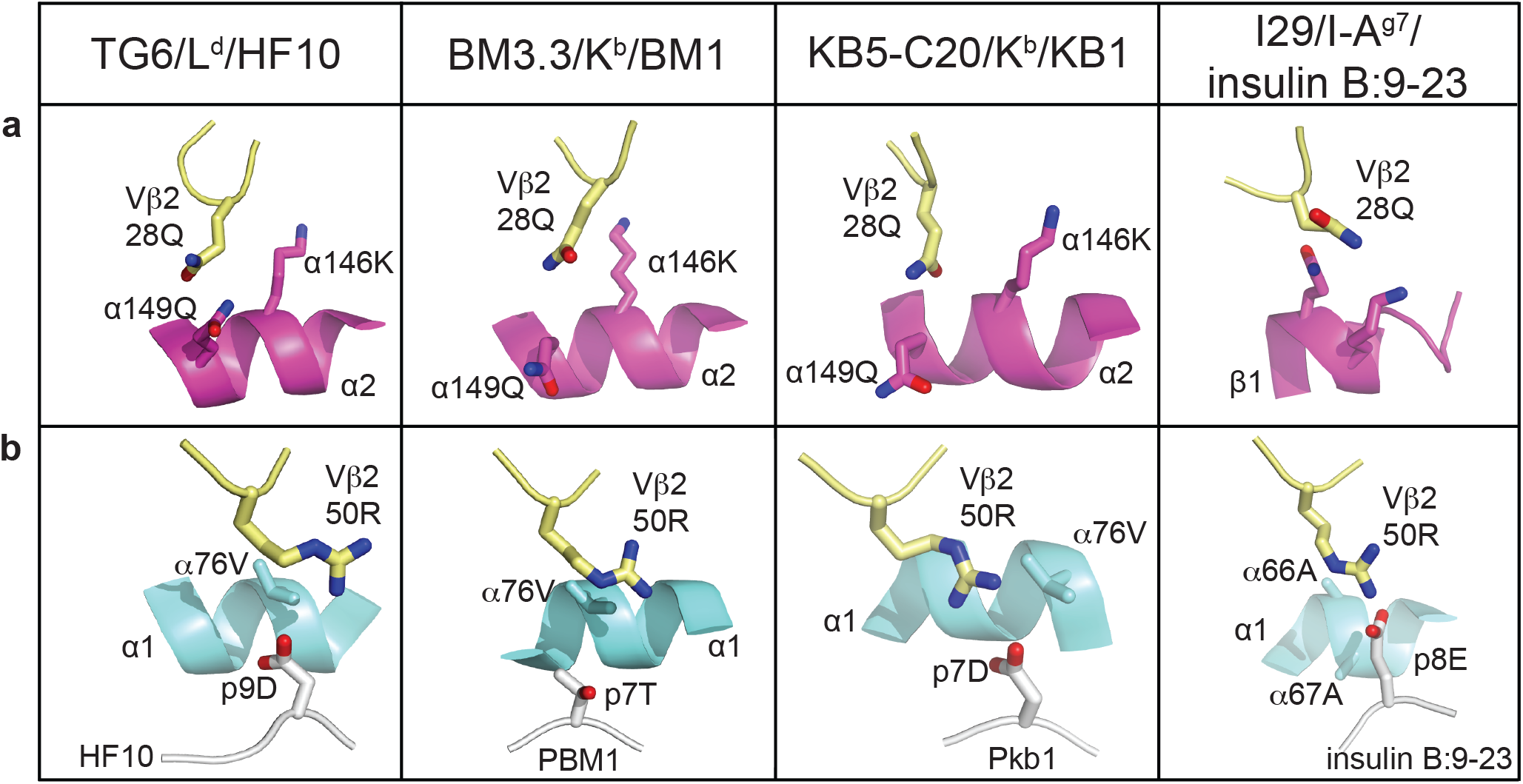
Conserved germline contacts of Vβ2 with pMHC. All Vβ2 TCR/pMHC structures are superposed and presented in the same view. Atoms are shown in CPK coloring. Vβ2 residues (28Q and 50R) are shown as pale yellow sticks. **a**, The position of Vβ2 28Q from Vβ2 containing TCRs is shown interacting with the MHC α2 helices (or β1 helix from MHC-II) shown in magenta ribbon diagram. **b**, The position of Vβ2 50R with pMHC contact from the same structures as in (a). MHC α1 helices are shown in cyan ribbon diagram, peptides are shown in white cartoon and the residues that interact with TCR are shown as white sticks. Protein Data Bank identifiers are: L^d^-HF10 (PDB: 6X31); BM3.3 TCR with K^b^-pBM1 (PDB: 1FO0); KB5 TCR with K^b^-Pkb1 (PDB: 1JK2); I29 with IA^g7^-insulin B:9-23 (PDB: 5JZ4).

## Discussion

It has been proposed that conserved germline contacts between TCR CDR1/2 residues and MHC help to impose the characteristic diagonal footprint of TCR on pMHC, although this remains controversial^2, 3, 4^. While investigating the structural basis of a potent CD8 T cell response to the parasite antigen HF10, we noted that the L^d^-HF10 specific TCR TG6, as well as other Vβ2-containing TCRs, adopt a parallel footprint on pMHC due to an unusual Vα-Vβ domain interface and TCRβ CDR1 and 2 loop conformations. This parallel footprint corresponds to a distinct set of conserved Vβ2-specific germline contacts, including one between CDR2 and the peptide. Thus Vβ2 represents “the exception that proves the rule” and solidifies the concept that conserved germline-encoded contacts help to define the binding orientation of a TCR on its pMHC ligand.

The distinct TCR footprint described here is observed with all four reported Vβ2 containing TCR-pMHC structures, regardless of the Vα chain (TRAV6D-7, 16, 14-1, 10) or MHC molecule (L^d^, K^b^, I-A^g7^), implying that the TCR Vβ usage has a major impact on the footprint of TCR on pMHC. Although mammalian genomes encode >30 functional TCRβ gene segments, they are not all equally represented in the mature TCR repertoire. In particular, Vβ8 is encoded by 3 related gene segments (TRBV13.1, 2, 3), but is expressed on ~50% of all T cells in mice, and represents >70% of published TCR-pMHC structures (Supplementary Fig. 4). Indeed, much of the evidence for germline-encoded TCR-MHC contacts comes from analyses of Vβ8-containing TCRs and structurally related Vβs in humans (Vβ13, 6, 7, 8 encoded by TRBV6, 7, 4, and 12 subfamilies respectively)^2, 5, 6^. While Vβ2 containing TCR-pMHC complex structures were reported earlier^26, 27, 28, 29, 30^ the unusual footprints on pMHC went unnoticed, in part because docking angles were calculated based on the positions of Vα and Vβ cysteines, an approach that does not capture the alterations in the footprint due to the unusual conformation of the Vβ2 CDR1 and 2 loops. With the addition of the TG6/HF10/L^d^ structure, Vβ2 is now the 2nd most well-represented Vβ segment amongst non-redundant mouse TCR-pMHC structure in the Protein Data Bank (https://www.rcsb.org/), making it possible to identify novel germline-encoded Vβ-specific contacts with MHC. While we were unable to identify an obvious counterpart to mouse Vβ2 amongst reported human TCR structures (data not shown), it seems likely that additional Vβ-specific pMHC docking patterns will emerge as the structures of additional TCR Vβ family-containing TCRs are determined.

In previous studies, the most prominent germline contacts involve tyrosine residues (e.g. Vβ8 Tyr 48 and Vα Tyr 29), which form extensive van der Waals contacts with the MHC α helices that make up the sides of the peptide binding groove^5, 6, 31^. Tyrosine residues are also highly represented in antibody CDRs and it has been suggested that the hydrophobicity and geometric flexibility of these contacts provides for relatively broad specificity, allowing for interactions with many allelic forms of MHC^2, 32, 33, 34^. In contrast, one of the prominent germline contacts in Vβ2 is an arginine in CDR2 which, in addition to van der Waals contacts with conserved aliphatic residues in the MHC α-helix, also forms a salt bridge with an acidic residue in the bound peptide in three out of four of the existing structures. This peptide-centric Vβ2 repertoire extends the conserved germline-encoded MHC interactions to a new V element. It is tempting to speculate that Vβ8 and Vβ2 containing TCRs may fill different niches in the immune response, with Vβ8 representing “generalists”, with a relatively broad specificity, and Vβ2 representing “specialists”, optimized for binding particular pMHC complexes (Supplemental Fig. 6). While TCR Vβ generalists would provide adequate responses and reliable coverage for many different pathogens, Vβ specialists could allow for “jackpot” responses that provide strong protection for particular pathogens. This is in line with the motif-driven, focused repertoire of L^d^-HF10 specific T cells reported here.

The notion of generalist versus specialist may also be applicable to the MHC-I molecule L^d^, which presents the HF10 peptide (Supplemental Fig. 6). In contrast to the broad peptide binding exhibited by most MHC-I molecules, L^d^ forms highly specific interactions with the antigenic HF10 peptide with bends in the peptide stabilized by multiple non-anchor residue contacts. Moreover six of the side chains are contacted by both the MHC and TCR, such that the specificity for peptide is shared between the MHC and the TCR. The highly specific binding between HF10 and L^d^ is consistent with earlier studies indicating that L^d^ has a constrained peptide binding site, binds poorly to self-peptides, and requires particular antigenic peptides to stabilize its cell surface expression^10, 19, 20^. Interestingly, human MHC alleles associated with elite control of HIV share polymorphisms with L^d^ that contribute to constrained peptide binding^7, 10^ and are also predicted to bind poorly to self-peptides^8^. Thus L^d^, as well as certain human MHC alleles associated with HIV control, may represent specialist MHC molecules which sacrifice broad coverage for the potential to generate highly protective T cell responses to particular peptides^15^.

The ability of MHC molecules to bind broadly to self-peptides has potential implications for how germline-encoded TCR-MHC reactivity is utilized in the mature TCR repertoire. It has been proposed that unfavorable interactions between CDR3 and self-peptides may counteract germline encoded reactivity to MHC to allow TCRs to avoid negative selection^2, 6^. In support of this idea, a TCR that was selected by a single peptide MHC complex, and therefore not subject to negative selection by diverse self-peptides, displayed exaggerated germline reactivity for MHC, leading to a high degree of cross-reactivity^6^. TCRs that are selected in the thymus by specialist MHC molecules may largely avoid the impact of negative selection due to lack of binding of self-peptides^8^. This may set the stage for jackpot T cell responses, since TCRs that recognize rare peptides that are able to bind to and stabilize the specialist MHC can take full advantage of both germline reactivity and peptide specificity to generate high affinity responses. This strategy may be particularly potent for Vβ2-containing TCRs, since specialist MHC molecules would be less likely to present self-peptides with acidic residues near the C-terminus, and thus would avoid strong self-reactivity leading to negative selection of thymocytes bearing these TCRs.

Previous efforts to understand T cell antigen recognition have largely focused on the most prevalent strategies, which may have evolved to provide broad and adequate coverage for most infections. In contrast, the current study highlights alternative “specialist” strategies for generating highly effective responses. Less commonly used Vβ gene segments and MHC alleles may provide a reservoir of recognition components that have the potential to provide highly focused and effective responses, and which could confer a selective advantage when populations are faced with particularly challenging pathogens. A better understanding of the recognition strategies used by specialist MHC alleles and Vβ segments should aid in the rational design of TCRs and improve our ability to target T cell responses in individual patients.

## Methods

### Animals

B6(C57BL/6) and B6.C (B6.C-H2d/bByJ) were obtained from The Jackson Laboratory (Bar Harbor, ME). TCR transgenic mice specific for L^d^-HF10 (TG6) were bred in house. Generation of TG6 mice was previously described^12^. In order to monitor multiple *T. gondii* epitopes, F1 mice (B6xB6.C) expressing both the H-2^b^ and H-2^d^ MHC class I molecules were used for all experiments. Six to ten week-old mice were used in all experiments. All mice were bred in UC Berkeley animal facility and were used within the approval of the Animal Care and Use Committee of the University of California.

### Infection

Mice were orally fed 70-80 cysts or injected intraperitoneally (i.p.) with 1×10^5^ live tachyzoites from the type II Prugniaud-tomato-OVA strain (Pru)^35^. This strain harbors immunogenic T cell epitopes derived from the parasite proteins, GRA6 (HF10 peptide)^11^, GRA4^36^, and ROP5^16^, and is engineered to express red fluorescent protein (RFP).

### Flow Cytometry

All antibodies were from eBioscience (San Diego, CA), Biolegend (San Diego, CA), or Tonbo (San Diego, CA). All tetramers were obtained from the NIH tetramer facility (Atlanta, GA). The tetramers were made by conjugating the biotin-labeled monomers with PE-labeled streptavidin (Prozyme, Hayward, CA) according to protocols from the NIH tetramer facility. All flow cytometry data were acquired by BD LSR Fortessa analyzers (BD Biosciences) and were analyzed with FlowJo software (Tree Star, Ashland, OR). Fluorescent AccuCheck counting beads (Invitrogen) were used to calculate total numbers of live lymphocytes.

### Single Cell TCR Sequencing

Mice were infected i.p. with the Pru strain of *T. gondii*. Spleens were harvested at 3 weeks post infection and L^d^-HF10 tetramer positive CD8 T cells were single-cell sorted into 96-well plates. TCRα and TCRβ sequences were obtained by reverse transcription and nested PCRs as described^37^.

### MHC I stabilization assay

RMA-S.L^d^ cells were obtained from N. Shastri (UC Berkeley). This assay was performed as previously described^17^. In brief, RMA-S.L^d^ cells were incubated at 37 °C, 5% CO2 for 8 h to saturate the culture medium with CO_2_, and then at room temperature overnight. The next day, cells were washed with PBS and plated at 3×10^5^ cells/well in a 96-W plate. Peptides of interest were added to the cells in serial dilutions. The plate was incubated for 1 h at RT and 3 h at 37°C. Cells were stained with the 30-5-7 antibody (specific for conformed, peptide-bound L^d^) and a goat anti-mouse IgG phycoerythrin (PE)-conjugated secondary antibody and analyzed by flow cytometry.

### Analysis of T cell activation

For analysis of the potency of HF10 peptide variants on TG6 T cell activation, RBC lysed splenocytes from TG6 TCR transgenic mice containing 10^5^ TG6 T cells were cultured in triplicate wells at 37 °C in 5% CO_2_. HF10 peptide variants were added to the cells in serial dilutions. Samples were harvested 48 hours later, stained for surface CD8, L^d^-HF10 tetramer, CD25 and CD44, and then analyzed by flow cytometry.

### Vβ usage of *T. gondii* epitope specific CD8 T cells

F1 (H2^bxd^) mice were infected with Pru strain *T. gondii* parasites. 3 weeks post infection, RBC lysed splenocytes were stained for surface CD8, peptide-MHC tetramer, CD44, and individual Vβs, and then analyzed by flow cytometry.

### Immunization with peptide-loaded dendritic cells

Bone marrow derived dendritic cells from male mice were incubated with 1μM of peptide for 3 hours at 37°C. Cells were washed and 5×10^6^ peptide-loaded dendritic cells were injected subcutaneously into naive F1 (H2^bxd^) female mice. Mice were sacrificed 7 days post immunization. Peptide sequences: HF10: HPGSVNEFDF; ROP5: YAVANYFFL; GRA4: SPMNGGYYM.

### Protein expression and purification

We used two systems to generate L^d^-HF10. For biophysical and crystallographic studies of pMHC, we produced soluble L^d^-HF10 by baculovirus infected insect cell expression^38^. The DNA encoding L^d^ (α1-α3) and HF10 (or alanine substituted variants) fused via a linker to β2m were cloned into pbac plasmid under polyhedrin and p10 promoters to produce secreted soluble Ld-HF10 from Hi5 insect cells. In this construct, L^d^ Tyr 84 and Gly at peptide p12 position from the linker that attaches β2m to pHF10 were mutated to cysteines to form a disulfide bond^39^. Secreted L^d^-HF10 in insect cell medium was captured with immunoaffinity chromatography and further purified by HPLC superdex 200 10/300 GL size exclusion column. For the crystallography of the TCR/peptide/MHC ternary complex, we produced a modified version of L^d^ variable region (α1-α2) bearing several mutations to improve stability^21, 40^. We restored arginine at L^d^ 97 back to Tryptophan as 97W is required for TG6 activation (data not shown). L^d^ (α1-α2) was expressed in *E. coli* BL21 as an inclusion body, solubilized in 8M urea, and refolded with synthetic HF10 peptide (HPGSVNEFDF, ordered from Peptide 2.0 Inc). Refolded L^d^-HF10 was further purified with HiLoad Superdex 200 26/600 size chromatography column.

TG6 α and β TCR chains were also produced by both baculovirus insect cell and bacterial expression systems. Acid-base leucine zipper stabilized, soluble TG6 molecules were produced in baculovirus infected Hi5 insect cells and enzymatically biotinylated for SPR study. For structural study, Vα and Vβ TG6 sequences were fused to the pET30 vector with human Cα chain as previously described^6^. The TG6 α and β TCR vectors were transformed separately into *E.coli* BL21. TG6 α and β proteins inclusion bodies were solubilized, mixed, and refolded by dialysis. The refolded TCR was further purified with a HiLoad Superdex 200 26/600 size chromatography column, followed by a Mono Q ion exchange chromatography.

### Surface plasmon resonance measurements

Approximately 2000 RU of biotinylated TG6 TCR was captured in the flow cells of a BIAcore streptavidin (SA) BIAsensor chip. Various concentrations of insect cell produced L^d^-HF10 and its mutated variant peptides were injected to the sensor chip and the association and dissociation kinetics were recorded and then corrected for the fluid phase SPR signal using the data from the biotinylated mouse BDC2.5 TCR. Kinetics was analyzed with BIAcore BIAEval 4 software.

### Protein crystallization

All crystals for data collection were produced by the hanging-drop vapor-diffusion method. Crystals of L^d^-HF10 were obtained at room temperature at a concentration of 7 mg/ml. The crystallization condition was 25% PEG 3350, 0.1M citrate pH5.5. TG6 TCR alone was crystallized at concentration of 10 mg/ml at 4°C in 20% w/v PEG10K, 0.1M Sodium citrate tribasic dihydrate pH5.5 and 1M Lithium sulfate monohydrate. For the TG6 / L^d^-HF10 complex crystallization, refolded L^d^-HF10 protein and TG6 TCR were mixed at 1:1 molar ratio at a final concentration of 10 mg/ml in 11 % w/v PEG 8K, 0.1 M MES 6.0, 0.24 M Ammonium sulfate.

### Data collection, data processing and structural analysis

All diffraction data sets were collected at synchrotron beamline ID-24C at the Advanced Photon Source, Argonne National Laboratory using the Pilatus detector. Initial models were solved by molecular replacement. Data collection and refinement statistics are shown in Table S1. The X-Ray diffraction data were collected under liquid-nitrogen cryo conditions at 100° K. The protein crystals were flashed-cooled in liquid nitrogen after a short soak in a cryo-protection solution consisting of the crystallization solution with 25% glycerol added. The data were indexed, integrated, scaled, and merged using the HKL2000 program^41^, the structures were solved by molecular replacement method using Phaser^42^ software and further refined by refmac5^43^, and rebuilding of the structure was performed by Coot^44^. NCONT in CCP4 was used to analyze the atom-to-atom contacts between the TCRs and pMHC^45^. Buried surface area (BSA) (Å2) is calculated with the PISA program from the CCP4 package^46^. Graphical representations of structures were constructed with PyMol (Schrodinger, LLC). The atomic coordinates and structure factors have been deposited in the Research Collaboratory for Structural Bioinformatics Protein Data Bank, https://www.rcsb.org (PDB ID codes are shown in Supplementary Table 1)

### TRangle determination and comparison

TRangle values of the TCR structures in the PDB were obtained from STCRDab on 26th April, 2019. New structures presented in this manuscript are annotated using the same pipeline as all structures in STCRdab. The TRangle was calculated using the protocol described in previous studies^25, 47^. In brief, the algorithm uses a defined set of the most structurally conserved positions in both the Vα and Vβ domains. It then fits reference frames through interface positions and computes the deviation from the pivot axis, C. The length of C is the dc distance. BA describes a torsion angle between the Vα and Vβ domains. BC1 and AC1 are angles that describe the tilt, while BC2 and AC2 capture the twist, between the two domains (Fig. 4c).

## Acknowledgements

We are grateful to staff at the 24-ID-E beamline at the Advanced Photon Source for assistance in synchrotron data collection. This work was supported by NIH Grants 5T32-AI-074491 (to Y.W.), ES-025797 and ES025885 (to S.D.), and RO1AI065537 and AI093132 to (E.A.R.), and the California Cancer Research Coordinating Committee (to A.T.).

## Author Contributions

Y.W., A.T., W.G., H.C., E.R., and S.D. designed the research. Y.W., A.T., W.G., H.C., Y. Z., and W. L. conducted experiments. Y.W., A.T., E.R., and S.D. analyzed data. Y.W., A.T., E.R., and S.D. wrote the manuscript. J.S. provided essential reagents. D.N. assisted in the synchrotron x-ray data collection. W.W. and C.D. performed the TRangle analyses. P.T. performed the Tdist analyses. E.R. and S.D. supervised the work.

## Author Information

The authors declare no competing financial interests.

## Data availability

The refined coordinates and structure factors for the X-ray structures of L^d^-HF10, free TG6 TCR, and TG6/L^d^-HF10 have been deposited in the Protein Data Bank with PDB accession codes: 6X2T, 6X30, and 6X31.

## Notes

### Competing Interest Statement

The authors have declared no competing interest.

