## Supplementary Figures for "Novel Vβ specific germline contacts shape an elite controller T cell response"

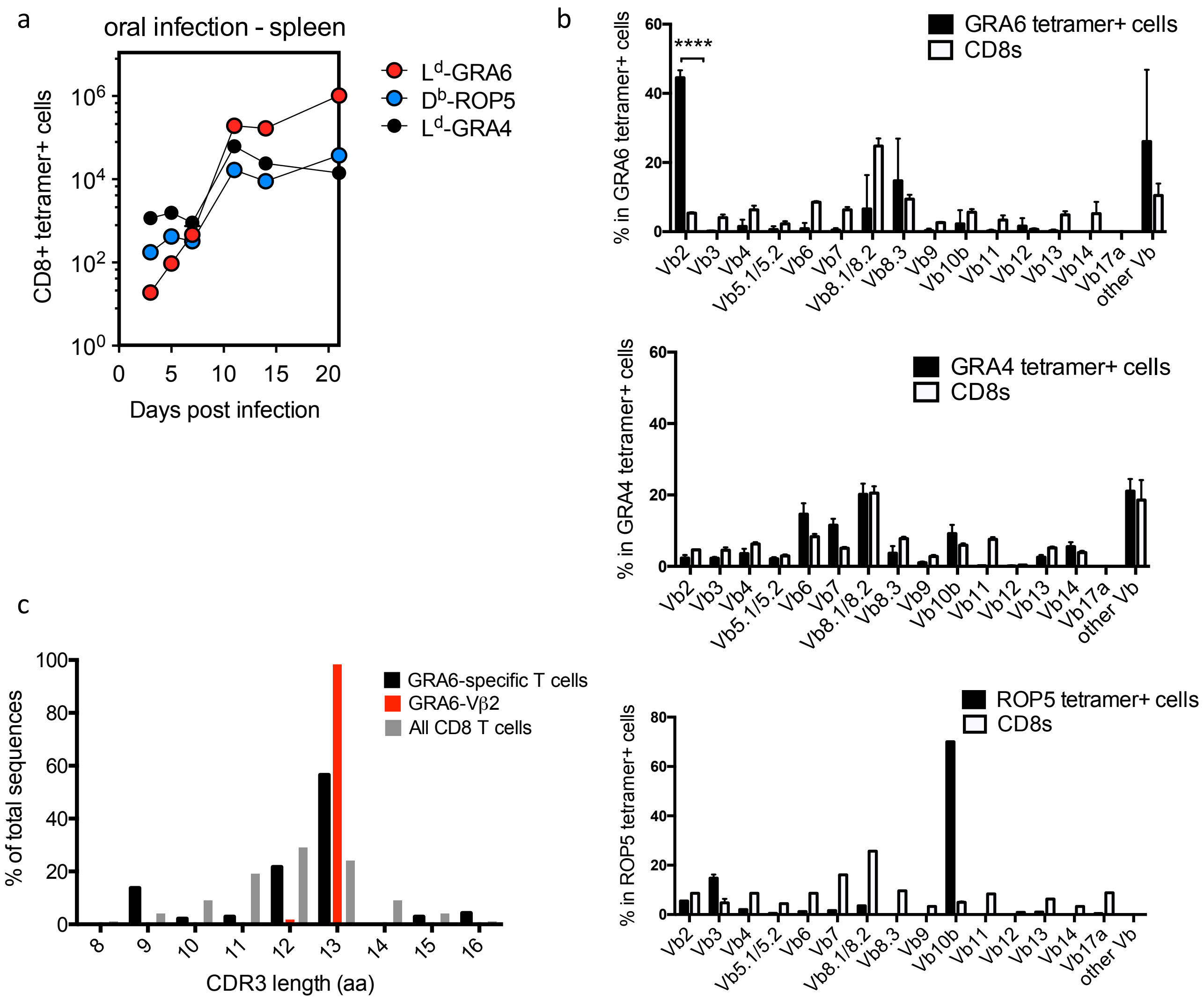


**Supplementary Figure 1 | Expansion, Vβ usage and clonal analysis of specific T cell populations after infection with *T. gondii* parasites.**

**a,** Mice were infected orally with cysts from the OVA expressing Pru strain *T. gondii*, and the numbers of the indicated pMHC tetramer$^+$ CD8 T cells were enumerated from the spleen using flow cytometry. **b,** Vβ usage of gated tetramer$^+$ splenic CD8 T cells was determined by flow cytometry. Vβ usage in total splenic CD8 T cells from the same mice is shown for comparison. Mice were infected i.p. with Pru strain *T. gondii* parasites and analyzed at 21 days post infection. Statistical significance was determined by a t-test (****$p<0.0001$). **c,** $L^d$-HF10 tetramer$^+$ CD8 T cells were single-cell sorted from mice 3 weeks after i.p. infection with Pru strain *T. gondii* parasites and TCRα and TCRβ genes from individual T cells were sequenced as described[36]. Restricted TCRβ CDR3 length in $L^d$-HF10 specific T cells from 2 individual mice. The Vβ2$^+$ $L^d$-HF10 specific T cells include 57 total sequences, from 22 distinct clones, of which 21/22 have a TCRβ CDR3 length of 13aa. The distribution of TCRβ CDR3 length in bulk CD8 T cells is provided for comparison.

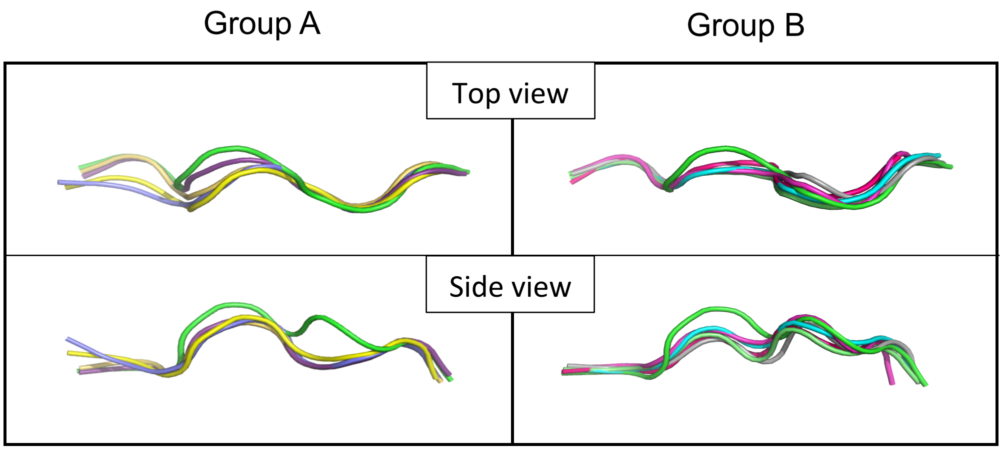


| | sequence | PDB |
|---|---|---|
| HF10 | HPGSVNEFDF | |
| group A | YPNVNIHNF | 1LD9 |
| | QLSPFPFDL | 1LDP |
| | SPAEAGFFL | 4MS8 |
| | QPAEGGFQ | 4MVB |
| | VPYMAEFGM | 5N5E |
| group B | QLSPFPFDL | 2E7L |
| | QLSDVPMDL | 3TFK |
| | SPAPRPLDL | 4MXQ |
| | MPAGRPWDL | 4N0C |

**Supplementary Figure 2 | Conformation of peptides bound to H2-L$^d$.**

Conformation of the bound HF10 peptide (green) in comparison to other L$^d$-bound peptides (various colors). Peptides with a bend at p5 (HF10 and 5 other 9mer peptides in various colors) are on the left (group A), and peptides with a bend at p6/7 (HF10 and 4 other 9mer peptides in various colors) are on the right (group B). Upper panels show a top down view, and lower panels show a side view.

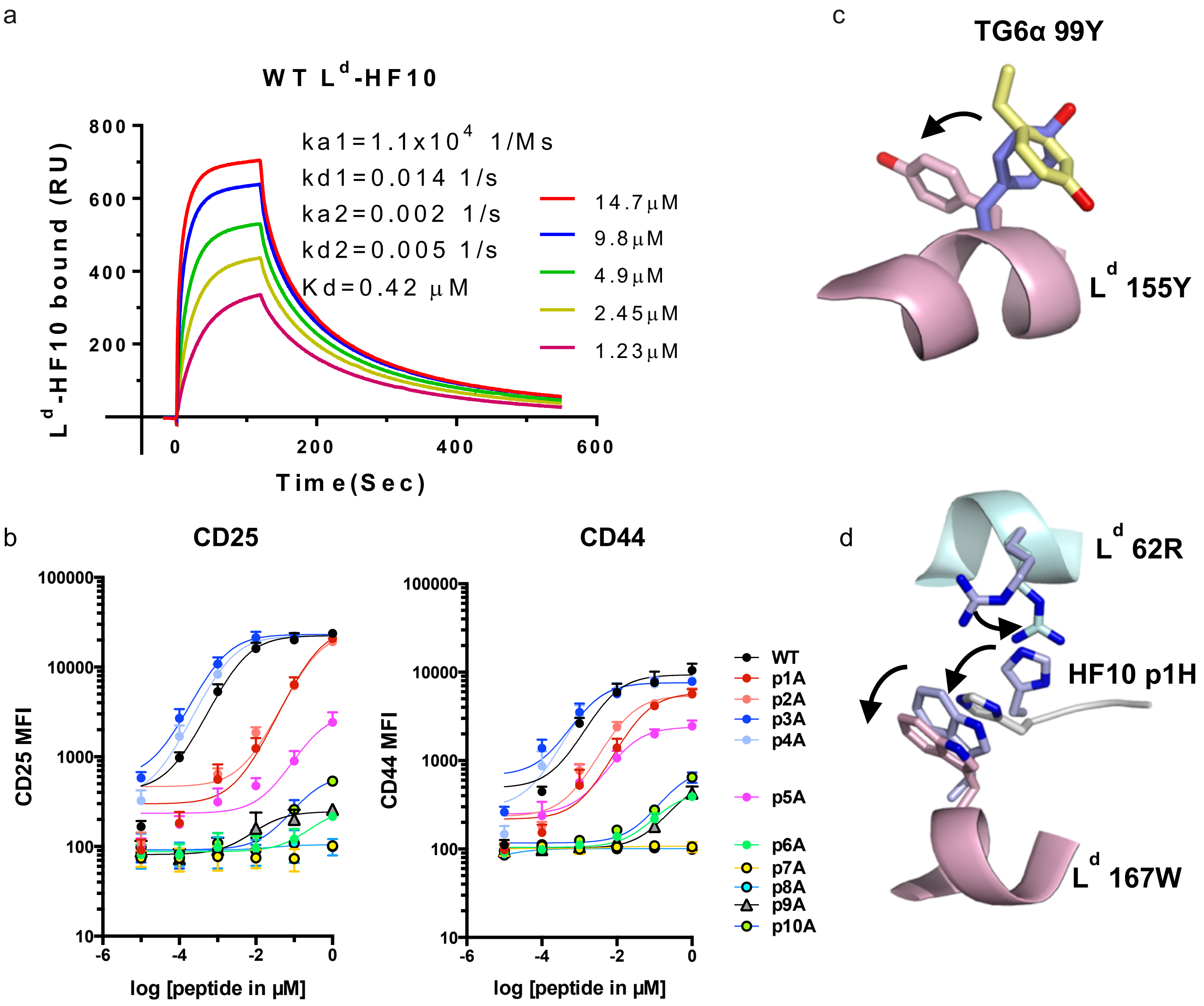


**Supplementary Figure 3 | Binding and activation of TG6 TCR.**

**a**, Corrected surface plasmon resonance (SPR) data for the binding of various concentrations of soluble L$^d$-HF10 to ~2000 RU of biotinylated TG6 TCR immobilized in flow cell on SA sensor chip. Another flow cell contained the irrelevant biotinylated mouse BDC2.5 TCR to correct for the fluid phase SPR signal. Standard BIA evaluation 4 software was used to calculate the kinetic and equilibrium constants. **b,** Naïve CD8 T cells from TG6 TCR transgenic mice were cultured with splenocytes incubated with WT HF10 peptide or the indicated alanine substituted variant peptides for 18 hours. Upregulation of surface CD25 and CD44 in response to TCR stimulation was measured by flow cytometry. **c,** Ribbon representation of a portion of L$^d$ α2 helix (light magenta) from the TG6/L$^d$-HF10 complex shown with side chains of 155Y. Superimposed is the side chain of 155Y prior to TG6 TCR engagement (blue) and the side chain of 99Y (yellow) from the TG6 Vα CDR3. **d**, L$^d$ α1 (light magenta) and α2 helix (pale cyan) with side chains of 62R and 167W (light cyan and light magenta), and cartoon representation of a portion of HF10 peptide with side chain of pP1H (silver). Superimposed are the side chains of L$^d$ 62R, 167W and p1H prior to TG6 TCR engagement (grey).

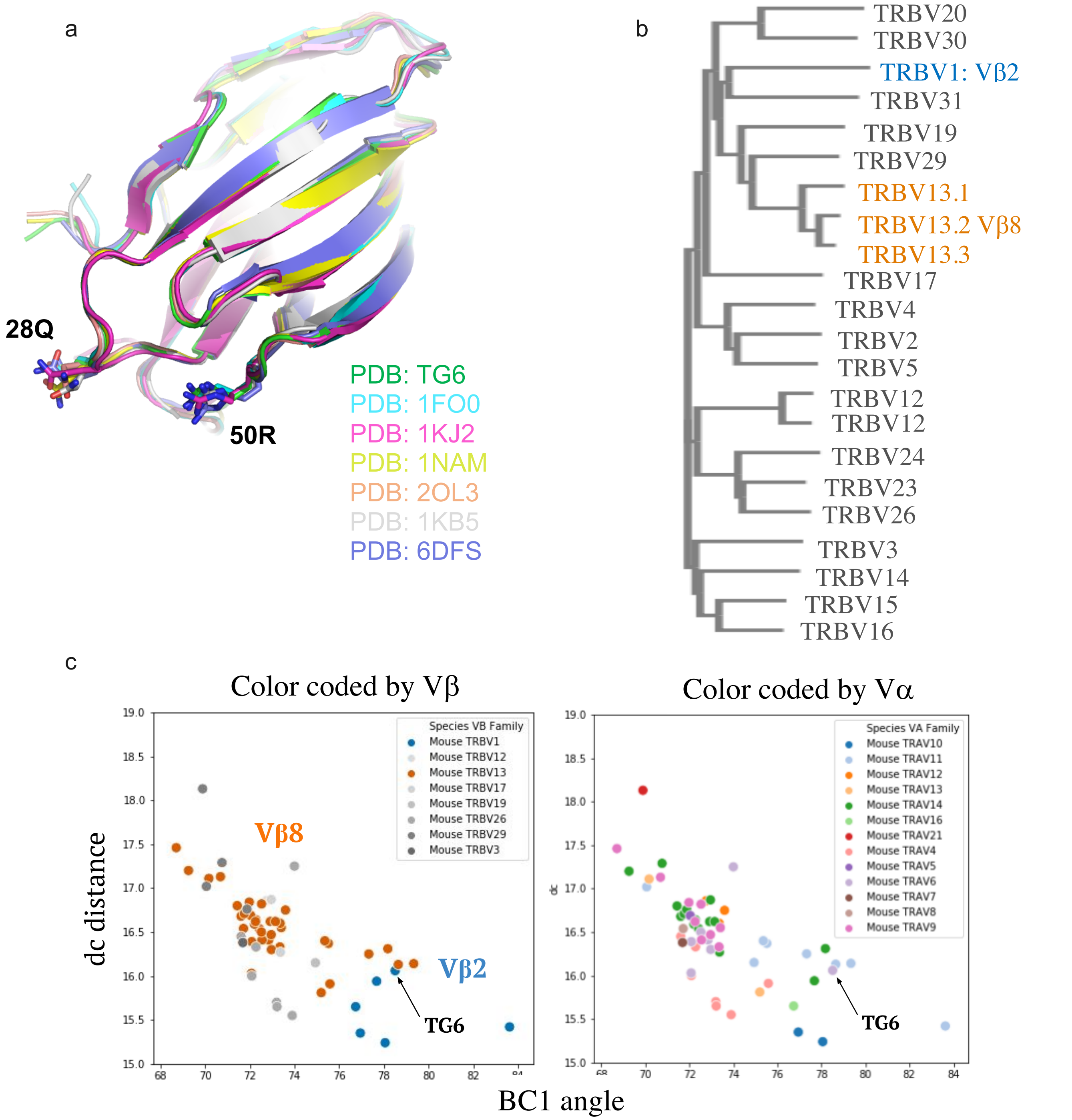


**Supplementary Figure 4 | Unusual CDR 1 and 2 loop conformation and Vα -Vβ interface geometry for Vβ2 containing TCRs.**

**a**, Ribbon diagrams of published Vβ2 containing TCR structures. The positions of conserved germline contact residues 28Q and 50R are shown. **b**, Tree showing relatedness of mouse TRBV genes based on inferred amino acid sequences. **c**, Plots of TRangle parameters dc distance versus BC1 angle for TG6 TCR (indicated by arrow) compared to non-redundant TCR structures in the PDB. Right panel shows data color coded to indicate TCR Vα usage. Left panel shows the same data color coded to indicate TCR Vβ usage and is reproduced from Fig. 4 for comparison.

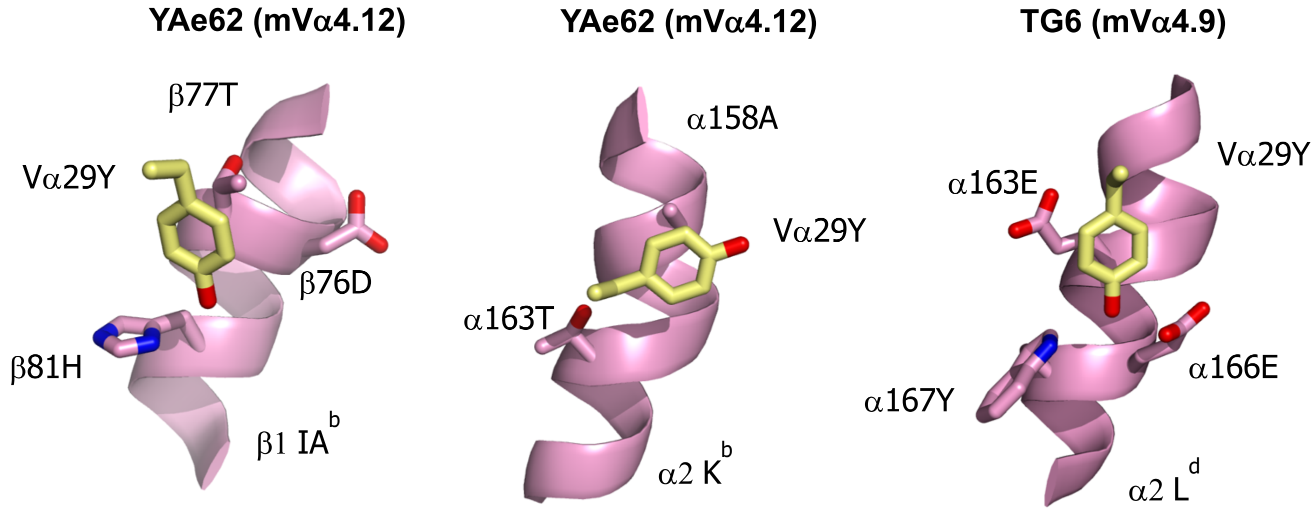


**Supplementary Figure 5 | Conserved germline contact between Vα CDR1 of TG6.**

MHC is shown in purple ribbon with selected side chains shown, and the TCRα germline contact residue 29Y shown in yellow. Atoms are shown in CPK coloring. Two published TCR structures are shown for comparison.

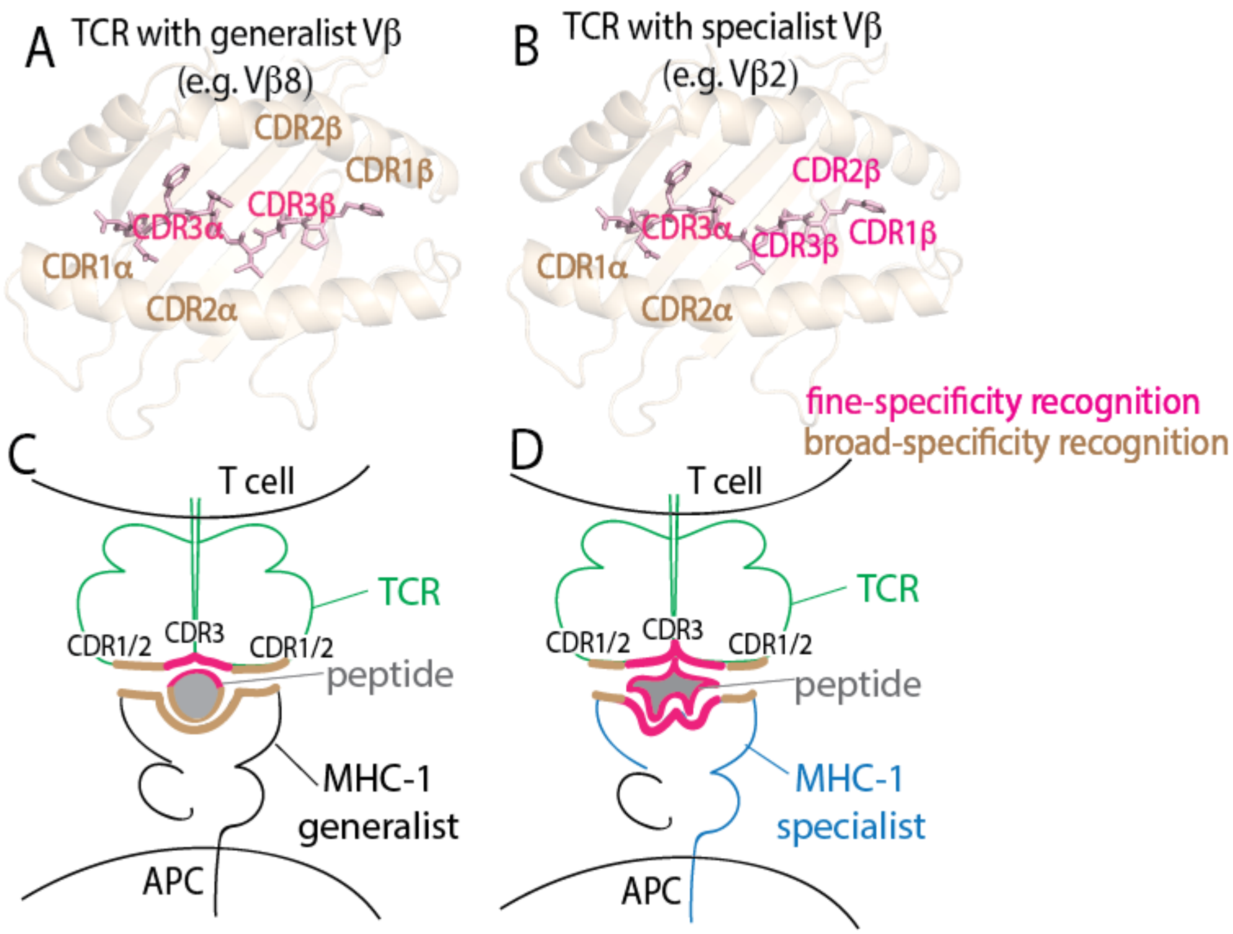


**Supplementary Figure 6 | Generalist and specialist strategies in T cell antigen recognition. a-b**, Interactions of TCR CDR loops with pMHC. MHC is shown as transparent brown ribbon diagram, and antigenic peptide is shown in pink stick, with positions of TCR CDR loops indicated. (a) For TCRs that contain generalist Vβ segments such as Vβ8, the highly variable CDR3 loops (purple text) are positioned over the peptide and provide for fine-specificity of ligand recognition, whereas the germline encoded CDR1 and 2 loops, (brown text), are positioned over the MHC, and provide for relatively broad recognition of different allelic forms of MHC. (b) For TCRs that contain specialist Vβ segments such as Vβ2, the TCRβ CDR loops are shifted away from the α1 helix of MHC, such that all 3 CDRβ loops can contribute to the fine-specificity for peptide. **c-d**, Diagram of TCR-peptide-MHC interactions with regions of broad specificity indicated by thick tan lines, and regions of fine specificity indicated by thick purple lines. (c) Generalist MHC-I molecules bind broadly to peptides, such that most of the fine specificity for peptides comes from the TCR. (d) Specialist MHC molecules such as MHC-I $L^d$ binding in a highly specific manner with peptide, such that the fine specificity for peptide is shared between the TCR and MHC.

**Supplementary Table 1 | Clonal analyses of TCR α and β chains of GRA6-specific T cells.**

| | TCRα | TCRβ |
|---|---|---|
| **TG6 sequence** | TRAV6-7/DV9, TRAJ52*01, CALGDPTGANTGKLTF | TRBV1, TRBJ2-1*01, CTCSAGRGGYAEQFF |
| Mouse 1 (1) | TRAV6-7/DV9, TRAJ30*01, CALRIQDTNAYKVIF | *TRBV1, TRBJ2-1*01, CTCSAGRGGYAEQFF |
| Mouse 1 (2) | TRAV6-7/DV9, TRAJ53*01, CALSEGNSGGSNYKLTF | *TRBV1, TRBJ2-1*01, CTCSAGRGGYAEQFF |
| Mouse 1 (3) | TRAV6-5, TRAJ32*01, CPGSSGNKLIF | *TRBV1, TRBJ2-1*01, CTCSAGRGGYAEQFF |
| Mouse 2 (1) | TRAV6-7/DV9, TRAJ52*01, CALSDRDSGGSNYKLTF | *TRBV1, TRBJ2-1*01, CTCSAGRGGYAEQFF |
| Mouse 2 (2) | *TRAV6-7/DV9, TRAJ52*01, CALGDPTGANTGKLTF | TRBV1, TRBJ2-1*01, CTCSAGRGAHAEQFF |

$L^d$-HF10 tetramer$^+$ CD8 T cells were single-cell sorted from mice 3 weeks after i.p. infection with Pru strain *T. gondii* parasites and TCRα and TCRβ genes from individual T cells were sequenced as described[36]. The TG6 TCRβ coding sequence from the original $L^d$-HF10 specific T cell hybridoma[8] was found independently in 2 additional mice, each time paired with a closely related TCRβ. Likewise, the TCR α gene of the TG6 TCR was found independently in one additional mouse, paired with a closely related TCRβ gene. (* indicates the identical TG6 sequence found in a new clone.)

**Supplementary Table 2 | Data Collection and Refinement Statistics**

| | H2-L$^d$-HF10 | TG6 | H2-L$^d$-HF10/TG6 complex |
|---|---|---|---|
| PDB code | 6X2T | 6X30 | 6X31 |
| **Data Collection** | | | |
| Space group | C2 | $P2_12_12$ | $P4_1$ |
| Cell dimensions | | | |
| *a*, *b*, *c* (Å) | 121.503<br>139.816<br>87.3441 | 86.16<br>196.331<br>63.112 | 91.168<br>91.168<br>106.247 |
| α, β, γ (°) | 90<br>130.824<br>90 | 90<br>90<br>90 | 90<br>90<br>90 |
| Resolution Range (Å) | 77-1.8 | 98-2.75 | 90-2.50 |
| $R_{sym}$ or $R_{merge}$*† | 0.09(0.53) | 0.01(0.1) | 0.03(0.29) |
| $I/\sigma I$* | 9.6(1.3) | 16(3) | 26.6(1.6) |
| Completeness (%)* | 99.6(90.1) | 98.5(90) | 99.8 (98.4) |
| Redundancy* | 1.9(1.8) | 1.2(1) | 6.4(5.5) |

*All data (outer shell).

† Rmerge = Σ(jI – 〈I〉 j)/Σ(I).

| | H2-$L^d$-HF10 | TG6 | H2-$L^d$-HF10/TG6 complex |
|---|---|---|---|
| **Refinement** | | | |
| Resolution Range (Å) | 50-1.8 | 50-2.75 | 5109-2.50 |
| No. reflections | 102152 | 26636 | 27675 |
| Rwork/ Rfree‡ | 19.91/22.66 | 23.89/29.45 | 17.16/21.79 |
| No. atoms | 6745 | 7152 | 5350 |
| Protein | 6267 | 7138 | 5118 |
| Ligand/ion | 98 | N/A | 210 |
| Water | 380 | 14 | 22 |
| B-factors | | | |
| Protein | 37.62 | 35.92 | 68.59 |
| Ligand/ion | 63.95 | N/A | 57.927 |
| Water | 40.25 | 15.60 | 80.55 |
| R.m.s deviations | | | |
| Bond lengths (Å) | 0.013 | 0.019 | 0.019 |
| Bond angles (º) | 1.664 | 1.934 | 1.932 |

‡ Rfree = ΣTestjjFobsj − |Fcalc|/ΣTest |Fobsj, where "Test" is a set of ~5% of the total reflections randomly chosen and set aside.

**Supplementary Table 3 | TG6 binding affinities to WT and mutants HF10 peptides**

| Peptide mutations | ka1 (1/M·s) | kd1 (1/s) | ka2 (1/s) | kd2 (1/s) | Kd (μM) |
|---|---|---|---|---|---|
| **HF10 WT** | $1.1X10^4$ | 0.014 | 0.002 | 0.005 | 0.42 |
| **p1A** | 5690 | 0.048 | 0.015 | 0.006 | 1.02 |
| **p2A** | 7980 | 0.066 | 0.009 | 0.014 | 1.84 |
| **p4A** | 1070 | 0.160 | 0.020 | 0.026 | 2.4 |
| **p5A** | 7510 | 0.284 | 0.009 | 0.014 | 1.85 |
| **p6A** | N/A | N/A | N/A | N/A | N/A |
| **p7A** | N/A | N/A | N/A | N/A | N/A |
| **p8A** | N/A | N/A | N/A | N/A | N/A |
| **p9A** | 52.7 | 0.175 | 0.008 | 0.009 | 170 |
| **p10A** | N/A | N/A | N/A | N/A | N/A |

Single HF10 residue alanine substitutions were tested to measure their effects on TG6 TCR binding.

**Supplementary Video Legends**

**Supplementary Video 1 | Rotation of the $L^d$-HF10 structure**

The $L^d$ $\alpha$1-$\alpha$3 sequence is shown as a silver ribbon diagram with the side chains of Trp97, Phe116, and Tyr156 shown. The bound $L^d$ peptide is shown as a green stick diagram with anchor residues Pro2 and Phe10 shown in yellow and the Glu7 shown in red. Note the bend in the peptide to accommodate the aromatic residues in the peptide binding site. Also note that the 10-mer peptide fits compactly within the peptide binding groove, with only the side chain of Glu7 pointing away from the MHC molecule.

**Supplementary Video 2 | Rotation of the TG6 TCR-$L^d$-HF10 structure**

The $L^d$ $\alpha$1-$\alpha$3 sequence is shown as a silver ribbon diagram, the HF10 peptide is shown as a green stick diagram with anchor residues Pro2 and Phe10 in yellow and the primary TCR contact residue Glu7 in red. The TG6 TCR $\alpha$ and $\beta$ chains are shown as red and tan ribbons respectively with the side chain of Arg97 shown in blue. Note the contact of Arg97 with Asp9 of the peptide and the close wrapping of the CDR3 loop around Glu7 of the peptide.

**Supplementary Video 3 | Comparison of the TG6 and Yae62 TCR structures**

The TG6 TCR (faint blue ribbon) is shown superimposed on the Yae62 (faint orange ribbon) with TCR$\alpha$ chains (very faint ribbons) aligned. The TCR$\beta$ CDR1 and 2 loops of both structures are in bold colors. Side chains of conserved germline contact residues are shown (Gln28 and Arg50 of TG6 TCR$\beta$ and Tyr48 of Yae62). Prolines 30 and 52 in CDR1 and 2 loops respectively of TG6 TCR$\beta$ are shown in dark blue. Note the displacement of the CDR1 and 2 loops between the 2 structures due to the distinct TCR$\alpha$-$\beta$ interface and CDR loop conformations.
